# Gibberellin enhances germination of *Nolana mollis* and *Heliotropium pycnophyllum* seeds

**DOI:** 10.1101/2025.01.10.632421

**Authors:** Liesbeth van den Brink, Melissa H Mai, Lorenz Henneberg, Margret Ecke, Rafaella Canessa

**Affiliations:** Plant Ecology Group, University of Tübingen, Germany; ECOBIOSIS, Departamento de Botánica, Facultad de Ciencias Naturales y Oceanográficas, Universidad de Concepción, Chile; Department of Organismic and Evolutionary Biology, Harvard University, USA; Center for Plant Molecular Biology, University of Tübingen, Germany; German Centre for Integrative Biodiversity Research (iDiv) Halle-Jena-Leipzig, Leipzig, Germany; Institute of Biology, Martin Luther University Halle-Wittenberg, Halle (Saale), Germany

**Keywords:** germination, gibberellin, *Heliotropium pycnophyllum*, *Nolana mollis*, seed dormancy

## Abstract

Desert plants often exhibit seed dormancy, which enables seeds to wait out unfavorable conditions and germinate when there is enough water for the seedling to establish securely. Here, we tested several dormancy-breaking mechanisms for *Nolana mollis* and *Heliotropium pycnophyllum*, including storage temperature, scarification, hormones, germination temperature, and light cycles during germination. For both species, gibberellin enhanced germination significantly. All other treatments, or combinations of treatments, had no significant effect. Our results can be applied to restoration efforts in arid areas, or in urban areas in arid climates, where the use of native species reduces watering necessities.

## Introduction

Many plant species living in warm arid environments, like the Atacama Desert, are adapted to survive harsh conditions. These adaptations include seed dormancy for up to several years, which enables the seeds to delay germination until adequate precipitation and temperature provide favorable conditions for growth (Finch-Savage & Leubner-Metzger, 2006). The conditions needed to break seed dormancy depend on the specific dormancy. For example, in ecosystems with marked seasons and where rain occurs only in winter, the seeds of some species must experience a “hot summer” and the colder temperatures of winter before they can germinate. This mechanism ensures that the seeds do not germinate during warm months, when drought stress would impair the growth of the seedling (Finch-Savage & Leubner-Metzger, 2006). Some seeds also need to experience a specific length of time in the dark, as this usually means that they are covered with soil, which they need to settle (Penfield, 2017). Many of the seeds in these environments have a thick seed coat that protects them from UV and evaporation, but also makes it hard for the seed to absorb water (Gutterman, 2000). These seeds often need to be “worn down” by scarring of some kind (e.g. by sand or insects), allowing water to penetrate the seed coat to initiate germination (Pezzani & Montaña, 2006). However, this puts the seed at risk of herbivory, as insects might eat the seed as soon as they have access to it (Schelin et al., 2004).

Some *Nolana* species have a non-deep physiological dormancy (i.e., requiring scarification to germinate) instead of a deep physical dormancy (i.e., requiring a specific time before germination) (Hepp et al. 2020). Previous work attempting to germinate seeds from *Nolana* species found that some species needed a cut at the funicular scar (Hepp et al. 2020) and that gibberellin enhanced the results (Cabrera et al, 2015; Douglas & Freyre, 2006; Hepp et al. 2020). Douglas (2007) found that the seed coat of *Nolana* species does not inhibit water uptake, and soaking could loosen the funicular germination plugs. This study also showed that gibberellin increased germination in some *Nolana* species and that seeds that were stored longer had higher germination rates.

In the case of *Heliotropium* species, germination also increased in *Heliotropium europaeum* when gibberellin or cold stratification was used (Aliloo & Darabenijad, 2013; Vaiga-Barbosa & Pérey-García, 2016). Aliloo & Darabenijad (2013) also showed that seeds germinated better after being stored longer than 12 months, or when potassium nitrate (KNO_3_) or cold stratification was applied. *Heliotropium europaeum* also germinated better in light, and poorly when buried (Humphries et al, 2018). Similarly, Chauhan & Johnson, 2008 indicated that *Heliotropium indicum* germinated poorly when buried, and was sensitive to water stress. However, *Heliotropium eichwaldi* germinated better in the dark, in combination with gibberellin (Srivastava, 1977).

Based on the above studies, gibberellin is suggested to influence the germination of both *Nolana* and *Heliotropium* species. Gibberellin is a versatile plant hormone responsible for plant growth, development and stress responses. For example, gibberellin is known to mitigate abiotic stresses by modulating plant processes, including germination (Shah et al, 2023). Here, we tested the effects of gibberellin and other environmental factors on the germination of *Nolana mollis* (Phil.) I.M. Johnst. and *Heliotropium pycnophyllum* Phil.

## Material and methods

Seeds were germinated using different regimes. We used a full factorial setup, resulting in 48 treatments per species (Fig. 1). To see if the seeds had a dormancy that could be broken by high summer temperatures, we placed 50% of the seeds at 40°C and 50% at room temperature (approximately 21°C) for 14 days in paper bags. After this, for each temperature, we tested if the seed coat needed to be physically scarred for the seeds to germinate. For that, 30% of the seeds were untreated, 30% of the seeds were scarred with sanding paper, and the last 30% were treated with acid. For the acid treatment, seeds were placed for 30 minutes in 0.5ml pure sulfuric acid (H_2_S0_4_) in 2ml Eppendorf tubes. After 30 minutes, the sulfuric acid was discarded, and 1.5ml demineralized water (H_2_O) was added to dilute the sulfuric acid that was still present in the Eppendorf tubes. After 10 minutes, the diluted sulfuric acid was discarded, and seeds were rinsed in 1.5ml of demineralized water. The next day, the remaining liquid was discarded. We further tested the effect of germination-regulating hormones on germination rates. We placed half of the seeds from each previous treatment overnight in Eppendorf tubes with 500ppm gibberellin (5 days for *N. mollis* and 4 days for *H. pycnophyllum*), while the other half was left untreated.

**Figure 1.**
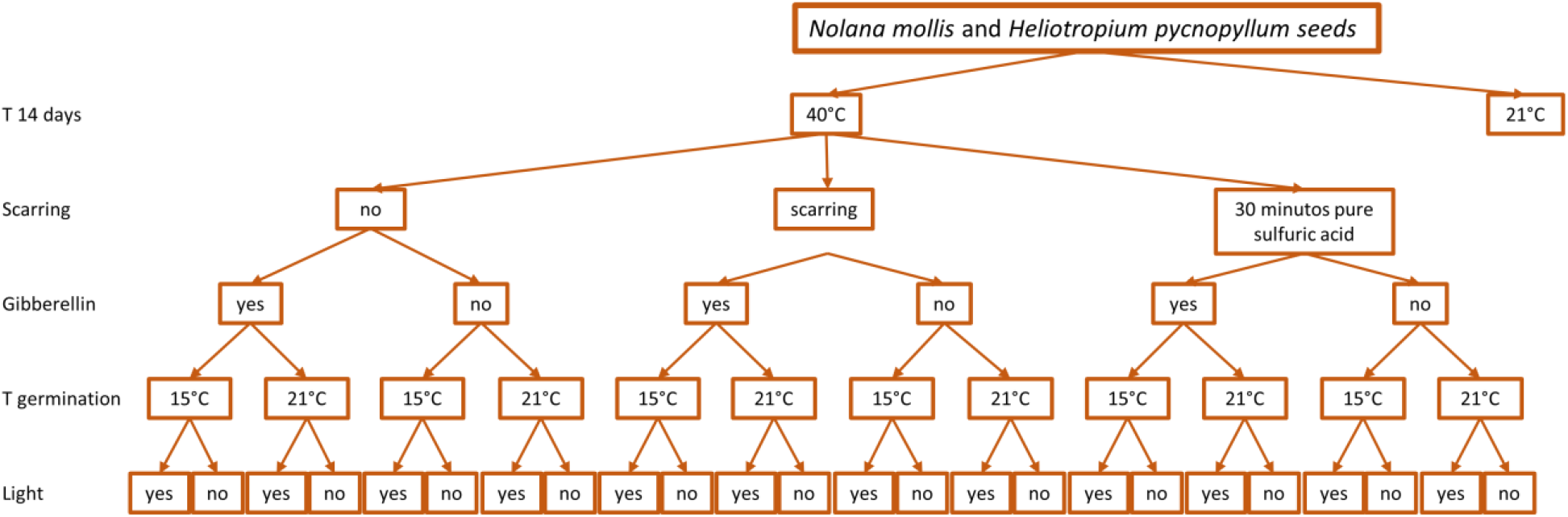
Setup of the experiment for the germination of seeds. The seeds were split into 48 different treatments. Note that only half of the treatments are depicted in the figure and that the scarification, gibberellin, germination temperature and light treatments were repeated for room temperature as well.

After the different treatments, all seeds were transferred to Petri dishes with a filter paper (4 discs per treatment, 48 in total, each with 10 seeds of *N. mollis* and 8 of *H. pycnophyllum*). We added a dental cotton roll and watered the cotton. The Petri dishes were then placed at a 45° angle, with the cotton roll at the bottom (Fig. 2). This ensured that the filter paper was moist but the seeds were not submerged. Half of the dishes (24 dishes) of each treatment were then placed at room temperature (21°C), of which half of the dishes (12 dishes) were placed in the dark and the other half without cover, thus experiencing daylight. The other half (24 discs) were placed at temperatures 15°C, and again half of these discs (12 discs) were placed in the dark and the other half without cover, thus experiencing daylight (Fig. 1). The plates were regularly watered and checked for germination.

**Figure 2.**
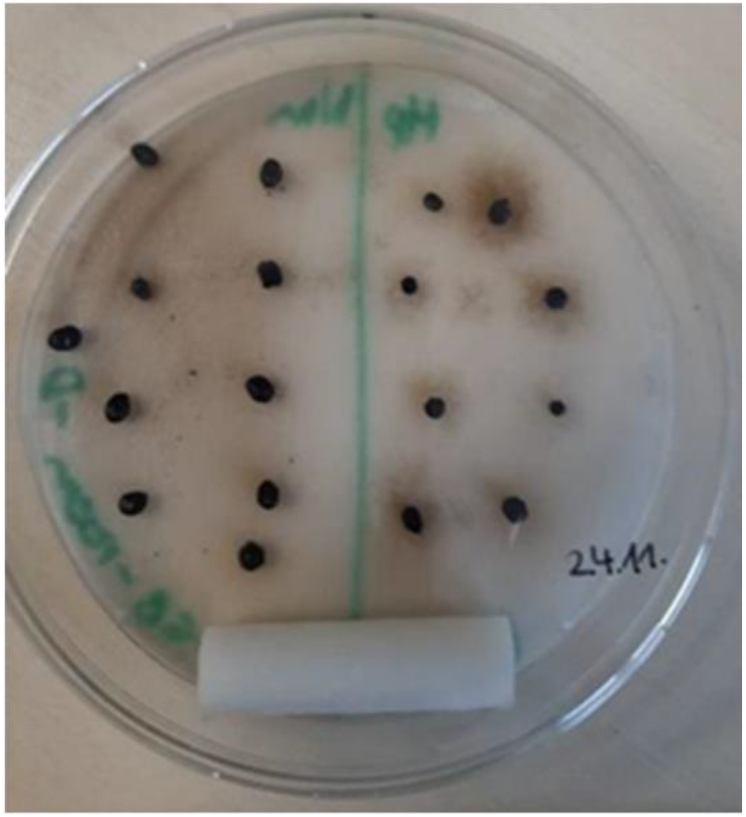
A Petri dish with ten *Nolana mollis* seeds (left) and eight *Heliotropium pycnophyllum* seeds on filter paper. At the bottom sits a cotton roll.

To test how the different treatments affected the germination we ran Chi-squared tests for each treatment. However, as the treatments were done in combination with other treatments, we also ran a multiple correspondence analysis to understand if the combination of some treatments would affect the germination as well.

We further investigated the germination rates of both species. The *N. mollis* germination experiments were distributed over four trials. Seeds were collected in March 2022 for the first two trials and in March 2023 for the latter two, and kept in the dark (in paper bags) at room temperature until the experiment started. In response to observed fungal growth on seeds from the first two groups, surface sterilization of the seeds was performed for the last two trials. To sterilize, the seeds were placed in 1.5 mL Eppendorf tubes and incubated for 1 min in 70% ethanol. The ethanol was then removed and replaced with 50% bleach (one part water, one part bleach with 6% NaOCl). After 5-10 minutes of incubation, the seeds were removed from the bleach and rinsed 5 times with deionized H_2_O that was sterilized through 0.2 μm nylon syringe filters (Whatman Cytiva 9910-1302) before incubating in 500ppm gibberellin (Sigma-Aldrich G7645) at room temperature for 5 days (*N. mollis*) or 4 days (*H. pycnophyllum*, as its seed coat is thinner). After the treatment, the seeds were placed in Petri discs, in a manner similar to the full factorial design described before. The seeds were regularly watered. This protocol was performed four separate times in October 2022, January 2023, July 2023, and October 2023 with a total of 1909 seeds. The unsealed plates were kept in a walk-in growth chamber with a 12h light (330uM) / 12h dark cycle at 21°C, 85-90% relative humidity. Germination was recorded at the first appearance of the radicle. Seeds were transferred to cells of 40% coarse sand and 60% potting mix when the root had grown to approximately 1-5 cm in length. Germination was determined as successful if it resulted in a viable seedling after planting. The resulting seedlings were kept in a walk-in growth chamber with a 12h light / 12h dark cycle at 21°C, with 60% relative humidity.

## Results

Gibberellin was the only treatment that significantly increased germination (p<0.001, Chi-squared tests) for both *N. mollis* and *H. pycnophyllum*). Storage temperature, scarring, germination temperature and light regime had no discernible effect on the germination (Fig. 3).

**Figure 3.**
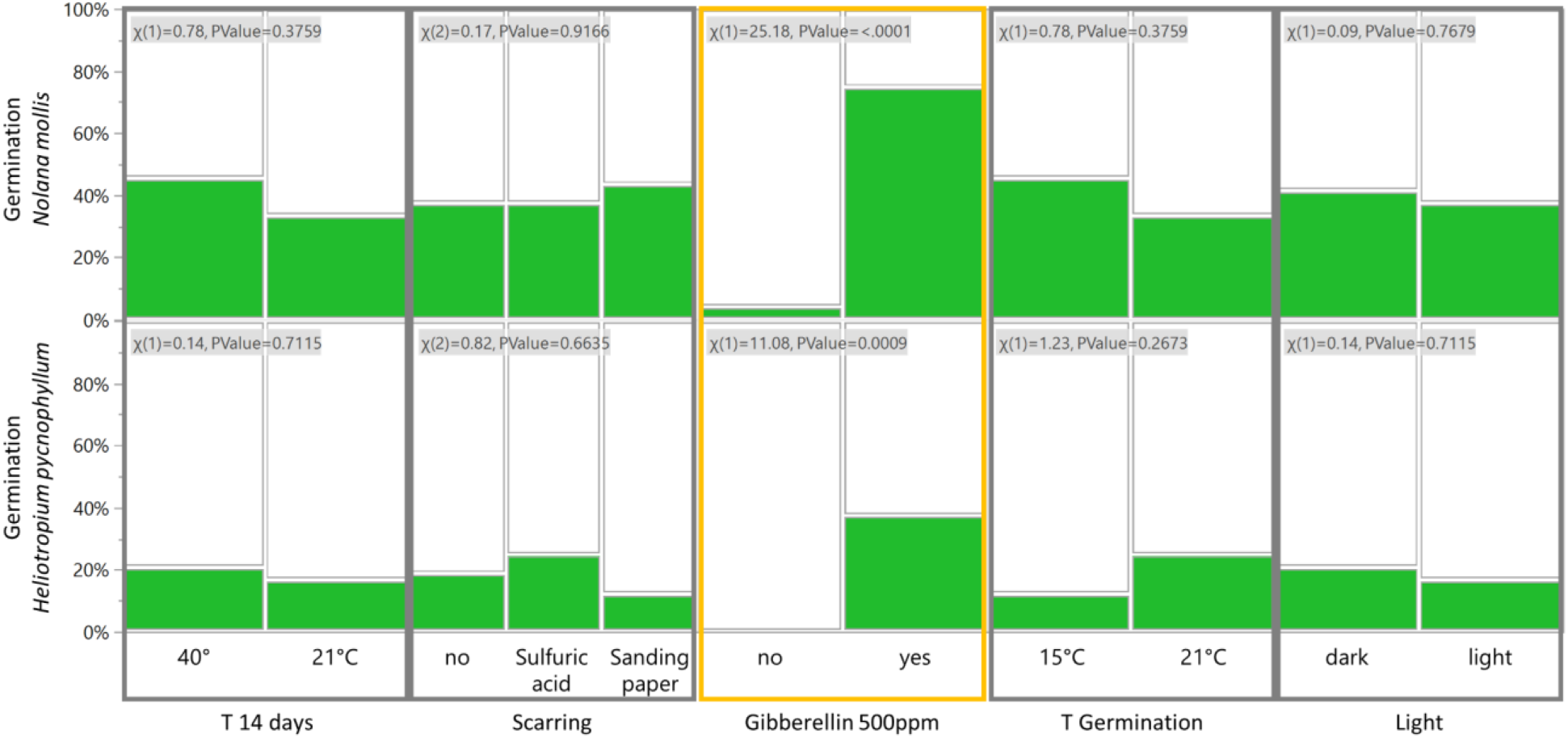
The effect of different treatments on the germination of *Nolana mollis* (top row) and *Heliotropium pycnophyllum* (bottom row).

The multiple correspondence analysis confirmed that gibberellin explained 93% of the germination of both *N. mollis* and *H. pycnophyllum* and that the other treatments had no effect (Fig. 4). Within 7–84 days, 58% of all seeds had germinated, half of which germinated within 28 days. However, only 5–20% of seeds in each trial resulted in viable plants (Figure 5), as several germinated seeds produced a radicle that arrested before reaching a length of 1 cm, and therefore would not have produced viable seedlings.

**Figure 4.**
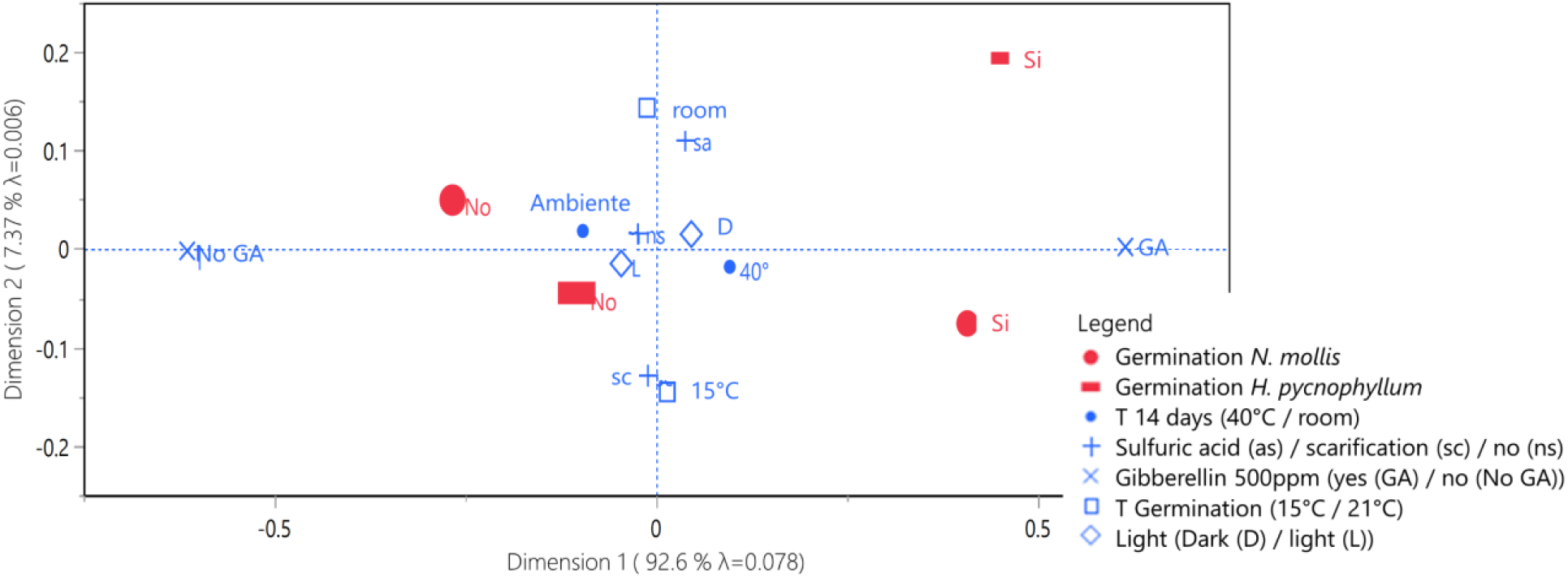
Effect size of treatments. Dimension 1 explains 93% and is dominated by the gibberellin treatment.

**Figure 5.**
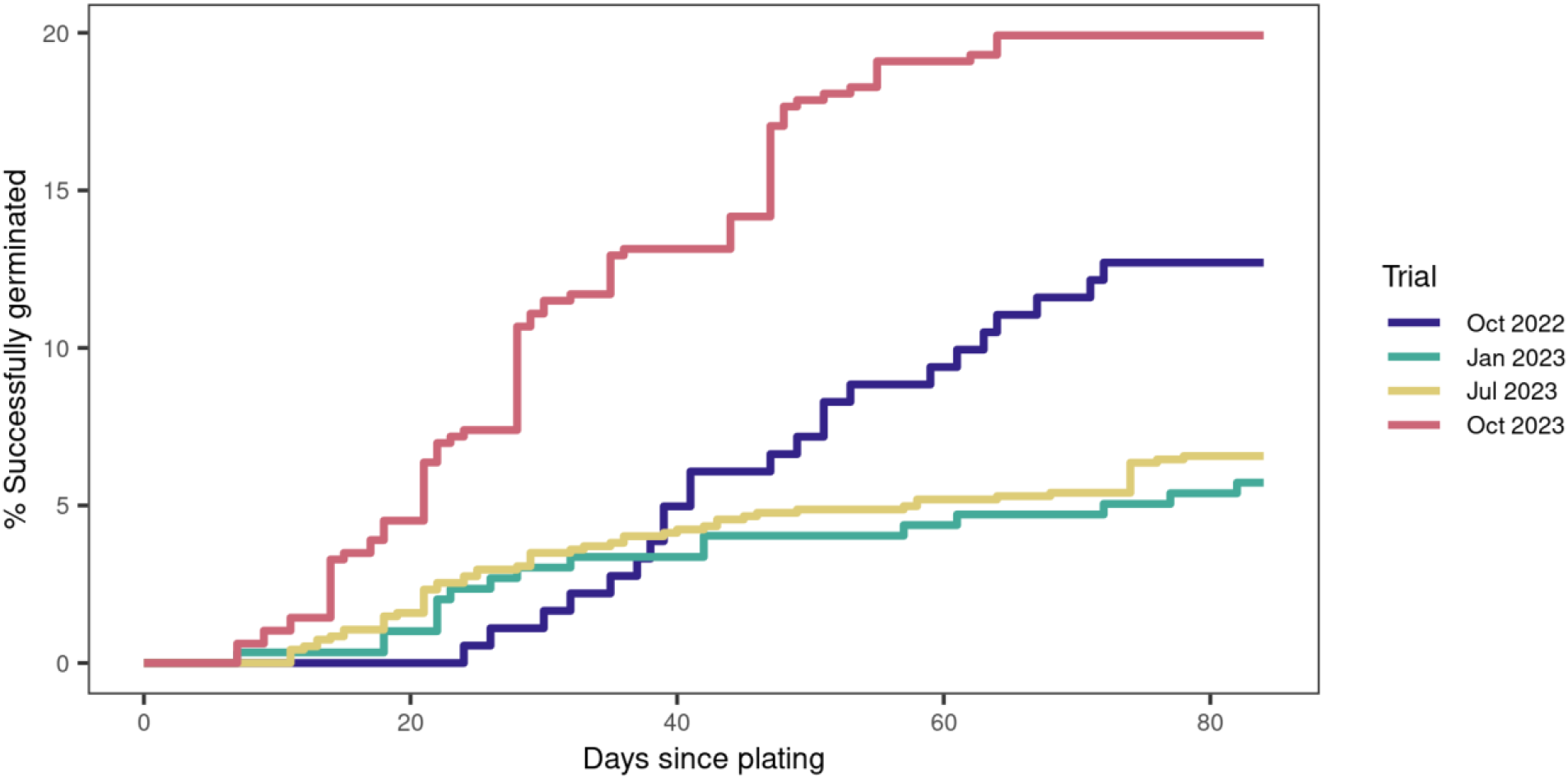
Cumulative germination rates for *Nolana mollis* as a percentage of the total number of seeds treated in each trial.

It was also observed that the *N. mollis* seed contains two funicular germination plugs and can occasionally give rise to two separate plants. Eleven seeds were observed to produce two seedlings per seed. These were identified by the emergence of two distinct shoots after potting or by the growth of a second radicle after the seed had detached from the cotyledons.

The first *H. pycnophyllum* seed germinated after 41 days. Forty-five days after the first germination, only 8% had germinated. It should be noted that most of these seedlings died when transplanted to pots.

## Discussion

The dormancy of *Nolana mollis* and *Heliotropium pycnophyllum* was not affected by high nor low temperatures. Moreover, neither scarring the seeds to break their dormancy nor exposure to dark conditions affected germination success. Instead, seeds of *N. mollis* and *H. pycnophyllum* germinated best after they had been treated with gibberellin (500ppm), a hormone that is responsible for several developmental processes, including germination (Shah et al, 2023).

Our experiment shows that it is possible to germinate seeds of *N. mollis* and *H. pycnophyllum* in controlled conditions like greenhouses and climate chambers. About 10% of the seeds successfully germinated, but it should be noted that we neither checked thoroughly for defects on the seeds nor performed any viability tests before starting the different treatments. As the seeds were collected from under the plants, some may have already been too damaged either by scarification (Pezzani & Montaña, 2006), microorganisms, or small insects (Schelin et al., 2004). Proper screening of seeds prior to treatment would likely improve the germination rate.

This germination experiment was part of a larger experiment in which the seedlings were transplanted to potted soil. It should be noted that most of the seedlings of *H. pycnophyllum* died after this transplant, making this a potentially unfavorable candidate species when planning experiments or restoration efforts. On the other hand, almost all seedlings of *N. mollis* survived the transplant to pots. This species is therefore a better candidate for restoration, as it requires fewer resources and experience to manage. Another *Nolana* species, *N. crassulifolia*, demonstrated even greater robustness in a parallel study by van den Brink et al. (which will be submitted to the technical journal *Biodiversidata*, volume 13, with the title “Comunicación breve sobre germinación de 15 especies nativas chilenas”). This species also needed 8 days until the first seeds germinated but, 51 days after the first germination, 20% of the seeds had germinated, and transplanting the seedlings was as easy as with *N. mollis*.

Our germination protocol can be used in greenhouses or laboratory conditions that cultivate native species, either to remediate environmental damage in the arid regions of Chile where these species are native, or for ornamental purposes in parks and plazas in northern Chile. The drought tolerance of these species makes them ideal for growing in the harsh environments of the towns in the Atacama Desert.

